# Understanding the selectivity of lysine negatively charged acyl modifications by OsCobB and its link with different stress conditions in plants

**DOI:** 10.1101/2025.04.25.649731

**Authors:** Asim Kumar Roy, Sonali Khan, Sanghamitra Dey

## Abstract

Recent proteomic studies have found several different lysine modifications in plants related to various important metabolic pathways. Lysine deacylation is a key feature of sirtuins for the regulation of several cellular processes in mammals. This may be involved in the tolerance against different stress conditions in case of plants. In this study, we deliver a comprehensive structural and kinetic analysis describing the selection of lysine acidic acyl groups by OsCobB. The catalytic efficiency (*k_cat_/K_m_*) of OsCobB to erase glutaryl, succinyl, and malonyl groups from the designated lysine site is favored due to the presence of Tyr55 and Arg58 at the active site. The selectivity of the substrate is further based on the OsCobB active site residues as well as residues next to the modified lysine. These modifications can be further linked to the plant’s mechanism to cope with stress due to dehydration, cold temperature and different metal toxicities.

## Introduction

Proteins undergo numerous post translational modifications (PTMs) in order to regulate various metabolic pathways and cellular processes [1]. Recently, researchers have expanded their horizon to study the diverse acyl modifications besides acetylation at the €NH_2_-lysine side chain [2–5]. Histone and non-histone proteins have been shown to have additional acyl-lysine modifications, such as malonylation (mal), succinylation (succ), glutarylation (glut), myristoylation, propionylation etc. [2,4,6] indicating their existence in living forms. The regulation of many of these newly discovered modifications is supported by the epigenetic players like sirtuins [3,4,7–9].

Sirtuins belong to a distinctively conserved class of NAD⁺-dependent protein deacylases and/or ADP ribosyltransferases. Majority of the studies have been done on humans, with some of them showing efficiency to remove the acidic lysine modifications. SIRT1, SIRT2 and SIRT3 are grouped together to display prominent deacetylase activity on its substrates [10,11] besides defatty-acylase effect. SIRT4, on the other hand, catalyzes weak ADP-ribosyl transfer [12] and lipoamidase activity [13]. SIRT5 has shown its favoritism towards succinylated, malonylated and glutarylated modifications, with its lower enzymatic efficiency for the acetylated substrates [2,3,7]. SIRT6 plays a crucial role in hydrolysing long chain fatty acids while its deacetylase activity is restricted to H3K9 and H3K56 [14,15]. On the contrary, SIRT7 was involved in H3 deacetylation at residue K18, causing transcription repression and in turn, controlling tumour growth [16]. Additionally, SIRT7 also showed detectable desuccinylation at H3K122 [17], Ezrin (EZR) [18], G6PD [19]and ADP-ribosyltransferase activities with PARP1, and transcription factors p53 and ELK4 [20–22]. So, out of all the seven sirtuins, SIRT5 and SIRT7 are known to remove the negatively charged lysine modifications. Proteomic studies have identified several substrates for lysine deacylation which are prevalent in nuclei as well as other cellular compartments including cytoplasm, chloroplast, mitochondria and plasma membrane. SIRT3, SIRT4 and SIRT5 are primarily localized in the mitochondria, with evidences of SIRT3 and 5 in cytosol and nucleus as well under various cellular conditions [9,23,24]. SIRT7 is localized in nucleoli with occurrences in nucleus [25].

PTMs have been repeatedly indicated to be critically important for plant’s adaptability to environmental stresses [26]. Mass spectrometric studies have shown the existence of lysine acylations on proteins with an expansive range of functions [27–29]. Mechanisms dealing with the addition or deletion of these acyl groups in plants have remained elusive for quite some time. Explorations of sirtuin family in plants classify them into three groups, namely II, III and IV. The annotation of acylations in rice proteome can be the stepping stone in deciphering its implications on physiological roles in plants. OsCobB was the first reported class III sirtuin, having high homology with bacterial CobB, and has a good deacetylase activity [30]. It does not have ADP ribosylase activity. It’s dual localization in the mitochondria as well as nucleus leave scope for the discovery of a huge array of functions in relation to deacylation. The other nuclear sirtuin, OsSRT1, is required to protect against cell damage in oxidative stress conditions [31]. It has mostly known deacetylation and ADP ribosylation activities [32]. Conversely, OsSRT2 is a mitochondrial protein with known decrotonylase activity [33]. In consideration of these findings, deacylase activities of OsCobB need to be explored with the intention of investigating its biological implications in rice. This would give an opportunity for future biological studies of these deacylations in plant physiology and functioning.

In our previous study on OsCobB, we have seen that this enzyme can interact with histones, H3 and H4 but not with H2A and H2B. Co-immunoprecipitation studies have shown its interactions with mitochondrial ACS and IDH2. OsCobB can effectively deacetylate them [34]. This current report laid down a detailed structural and kinetic analysis portraying the interconnection between OsCobB and its substrate acyl-group selectivity. Various acylated substrates are known to form intermediates in various important metabolic pathways like the TCA cycle, glycolysis and fatty acid synthesis in plants. MD simulation studies were carried out to investigate the dynamic behavior of these acidic lysine modifications while binding at the active site. Further, we investigated the catalytic abilities of OsCobB, correlating it with different biotic and abiotic stresses. It suggests that these reactions may be physiologically relevant affirming future research scope.

## Materials and methods

All the reagents used in this study were purchased from HiMedia laboratory, India, Sisco Research Laboratories (SRL) India and Sigma-Aldrich, USA. Antibody for ACS2 (ab66038), IDH2 (PHY0040S), acetyl Lys (9681S), Pan-malonyl (PTM-901), Pan-succinyl (PTM-419), Pan-glutaryl (PTM-1152) antibody were purchased. H3K9Ac peptide (ARTKQTARK(Ac)STGG), ACS peptide (KTRSGK(Succ)IMRRI) were synthetically prepared from Bio Basic Inc, Canada. Peptides corresponding to residues 4−17 of histone H3 (KQTARKacylSTGGKAPR) were synthesized from abclonal. Custom made antibody for full length untagged OsCobB was prepared from rabbit (Biobharati Lifesciences Pvt. Ltd, India).

### Molecular docking studies (*in-silico* studies)

Molecular docking studies were performed using HDOCK server (http://hdock.phys.hust.edu.cn/) to study the protein-ligand interactions at atomic level. OsCobB model was prepared from ITASSER server (https://seq2fun.dcmb.med.umich.edu/I-TASSER/) with good geometry. The chemical structures of the different acylated peptides (H3K9, H4K16, IDH2 and ACS) were drawn using PubChem sketcher. The best fitted docked conformations with good docking scores were used for further structural analysis. UCSF Chimera 1.16 software was used to visualize the docked complexes for the best orientation of the ligand with OsCobB and interaction studies [35]. LigPlot v2.2 (Laskowski and Swindells, 2011) was used to generate the 2-D schematic diagrams of protein-ligand interactions of the docked complexes. *In-silico* mutagenesis analyses were carried out using BeAtMuSiC prediction software (http://babylone.ulb.ac.be/beatmusic).

### *In-vitro* deacylation activity of OsCobB (dot blot method)

The recombinant OsCobB was purified using the protocol explained earlier [34]. H3K9 acylated peptides were synthesized and were used as substrates to identify the specificity of OsCobB towards the varied lysine acyl side chains. The reaction was performed at 25°C for 1-3 hrs, depending on the catalytic efficiency of OsCobB towards the modification. The reaction buffer was 50mM Tris pH 7.5, 150mM NaCl and 5mM DTT. A 20μl reaction was prepared using 0.8µM OsCobB, 1µg peptide and a gradient of 0-700µM NAD^+^. Then, dot blot analysis was done in triplicates. Different acyl modification-specific primary antibody (1:1000) was used to detect the intensity of dots (enzyme reactions). HRP-conjugated goat anti-rabbit (1:5000) is the secondary antibody.

### Deacylation activity of OsCobB with leaf extracts

Steady state kinetic analysis of OsCobB deacylation was performed by keeping the fixed concentration of the nuclear/mitochondrial extract and a varied concentration of NAD^+^ (0–600 μM). Reaction mix containing recombinant OsCobB (0.8 μM), NAD^+^, substrate protein (extracted from rice leaves (20μg)) in buffer (50 mM Tris (pH 7.5), 150 mM NaCl, 5 mM DTT) was incubated at 25°C for 1-3 hr. The reaction mix was resolved in 12% SDS-PAGE and was electro-transferred on nitrocellulose membrane. The blots were blocked using 5% skimmed milk + Tris based saline (TBS) for 1 h, followed by TBST buffer wash and incubating in acylation-specific primary antibody (1:1000) for 2h. This was followed by incubation with an HRP-conjugated Goat secondary antibody (1:5000) for 45min and developed with ECL reagent (Bio-Rad) using the Bio-Rad ChemiDoc MP imaging system. Protein band intensity was quantified in ImageJ. Steady state kinetic parameters were calculated using Non-Linear regression (Michaelis Menten (MM) or *k_cat_* Model) in GraphPad Prism version 9.1.2 (San Diego, USA).

### Comparison of extent of acylation with OsCobB expression during stress conditions

The amounts of the probable nuclear and mitochondrial substrates were compared in different stress conditions along with the expression level of OsCobB protein in the extracts. For this, nuclear and mitochondrial extracts were prepared as mentioned in [34]. 30μg nuclear/mitochondrial extract of normal as well as stressed varieties of rice leaves was loaded onto a 12% SDS-PAGE and western blot analysis was done. The degree of OsCobB expression was determined using anti-OsCobB primary antibody (1:1000). The extent of acylation was determined using anti-acyl lysine antibodies. The amounts of H3, H4, ACS and IDH2 was further determined using the protein specific primary antibody. The data was normalized using an external control (His-GST). All experiments were done in triplicates. The protein band intensities were measured using ImageJ software and bar graph was prepared by plotting the fold change in OsCobB expression in correspondence to the extent of deacylation in different stress conditions.

## Results and discussion

### An understanding of the catalytic acylations preferred by OsCobB

The ability of class III sirtuins to deal with diverse acyl modifications of their substrates has an impact on their physiological functions. Hence, understanding the selectivity of acyl modifications of different target molecules, binding at the active site of OsCobB would provide an idea about its biological roles in plants. Since OsCobB was previously reported to effectively deacetylate histone H3 at K9 position, this peptide was selected to maintain a consistency of docking and nullifying the effect of the peptide backbone on docking [30]. H3K9 peptides with the acidic modifications (KQTARK_Acyl_STGGKAPR) in lysine were further selected for this study. To analyse the effect of OsCobB towards various acyl groups, steady-state deacylation rates were compared after performing kinetics study using western blot. (**Fig. 1A, Table 1**). Further attempts were then made to correlate these *in-vitro* catalytic abilities of OsCobB with the *in-silico* findings.

**Figure 1:**
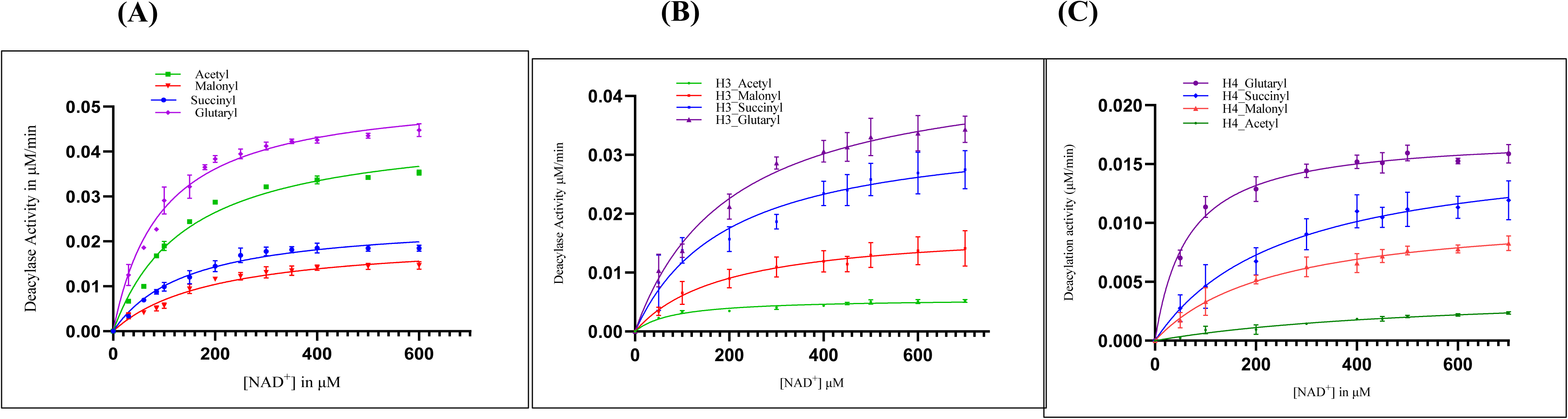
Michaelis-Menten plots comparing OsCobB-catalyzed deacylation activity of H3K9 peptide and endogenous histones H3 & H4. The MM plots of OsCobB for H3K9, overall histones H3 and H4 deacylations were calculated using Graphpad Prism 9.1.2 (MM Model). The error bar depicts the S.D.; n=3.

**TABLE 1:** A comparison of the kinetic parameters for NAD^+^ in OsCobB catalyzed deacylation of modified H3K9 peptides.

| **H3K9 acyl**<br>**peptide**<br>**modifications** | **$k_{\text{cat}}^{\text{app}}$ ($\text{min}^{-1}$)** | **$K_{\text{m}}^{\text{app}}$ for $\text{NAD}^+$**<br>**($\mu\text{M}$)** | **$k_{\text{cat}}^{\text{app}} / K_{\text{m}}^{\text{app}}$ ($\text{M}^{-1}\text{s}^{-1}$)** |
| --- | --- | --- | --- |
| Acetyl | $3 \pm 0.08$ | $144.5 \pm 11$ | 346 |
| Malonyl | $1.4 \pm 0.067$ | $198 \pm 23$ | 118 |
| Succinyl | $1.65 \pm 0.06$ | $145.6 \pm 16$ | 206 |
| Glutaryl | $3.6 \pm 0.07$ | $95.41 \pm 6.1$ | 652 |

OsCobB was capable of deglutarylate at H3K9 site with the highest catalytic efficiency in comparison to the other acidic peptide modifications, indicated by its higher *k_cat_/K_m_* ratio. The inclination of OsCobB towards the negatively charged acyl modifications can be justified as these modifications can penetrate deeper into the acyl-lysine cavity. Their chemical structures also show the addition of -CH₂ groups from malonyl to glutaryl (3-5 carbon), allowing the longer lengths of the modification to infiltrate deeper into the cavity (**Fig. 2A**). Acidic modifications are also known to change the charge of the lysine in a protein from +1 to −1 in comparison to 2-carbon acetyl group changing it to 0. 2-D representation of OsCobB docked structures strongly interacting with glutaryl (*k_cat_/K_m_* = 652 M^−1^s^−1^), succinyl (*k_cat_/K_m_* = 206 M^−1^s^−1^) and malonyl (*k_cat_/K_m_* = 118 M^−1^s^−1^) side chains owing to the YXXR motif (**Fig. 2B**). Human SIRT5, another class III sirtuin, was found to display a clear preference towards demalonylation and desuccinylation of its substrate, H3K9 in comparison to weak deacetylation (29 to >1000 fold) [3]. OsCobB shows comparable efficiency in erasing the acetyl as well as acidic lysine modifications at the K9 position.

**Figure 2:**
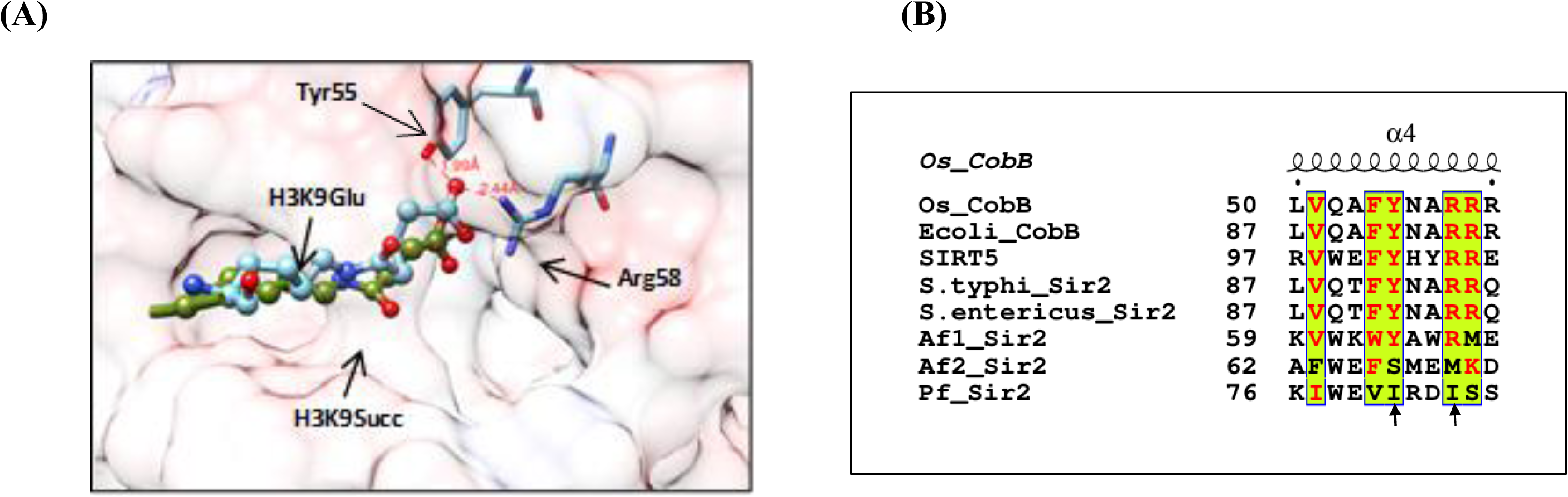
(A) Interaction of the modified lysine peptides at the substrate binding site of OsCobB (B) Conserved Tyr55 and Arg58 in class III sirtuin members in comparison to other members.

Despite *in-vitro* analysis showing a preference for acidic and acetyl modifications at H3K9 position, there is a possibility of OsCobB erasing the said modifications at other lysine sites. Also, there was a need to examine the behaviour of deacylase OsCobB with respect to full length histones H3 and H4. For this, we have extracted the histones from rice leaves. Even though H4 was not a good candidate for deacetylation in comparison to H3, there is a possibility that acidic modifications in H4 can be removed by OsCobB.

It can be noted that OsCobB is most efficient in desuccinylation, deglutarylation of histone H3, followed by H3 demalonylation (**Fig. 1B**). Comparing the extent of deacylation using histone H4 extracted from rice leaves demonstrated the dominance of OsCobB deglutarylase activity (*k_cat_/K_m_* =309 M^−1^s^−1^). In comparison to histone H3, acetyl, succinyl, malonyl modifications were not removed efficiently in H4 but OsCobB follows the same inclination of removal with increase in chain length. (**Table 2**). Overall, we can also say that H4 is not a good target for the removal of acetyl and acidic modifications by OsCobB.

**TABLE 2:** Kinetic studies to determine the preference of OsCobB towards H3 and H4 deacylations using nuclear extract of rice leaves.

| **H3 modifications**<br>**(Overall)** | **$k_{\text{cat}}^{\text{app}}$ ($\text{min}^{-1}$)** | **$K_{\text{m}}^{\text{app}}$ for $\text{NAD}^+$**<br>**($\mu\text{M}$)** | **$k_{\text{cat}}^{\text{app}} / K_{\text{m}}^{\text{app}}$**<br>**($\text{M}^{-1}\text{s}^{-1}$)** |
| --- | --- | --- | --- |
| Acetyl | $0.4 \pm 0.01$ | $83.2 \pm 11$ | 74 |
| Malonyl | $1.2 \pm 0.1$ | $190.4 \pm 48$ | 105 |
| Succinyl | $2.4 \pm 0.21$ | $209 \pm 54$ | 188 |
| Glutaryl | $3.0 \pm 0.24$ | $247 \pm 52$ | 202 |

| **H4 modifications**<br>**(Overall)** | **$k_{\text{cat}}^{\text{app}}$ ($\text{min}^{-1}$)** | **$K_{\text{m}}^{\text{app}}$ for $\text{NAD}^+$ ($\mu\text{M}$)** | **$k_{\text{cat}}^{\text{app}} / K_{\text{m}}^{\text{app}}$**<br>**($\text{M}^{-1}\text{s}^{-1}$)** |
| --- | --- | --- | --- |
| Acetyl | $0.3 \pm 0.05$ | $574 \pm 167$ | 8.7 |
| Malonyl | $0.73 \pm 0.04$ | $235 \pm 37$ | 52 |
| Succinyl | $1.1 \pm 0.1$ | $257 \pm 61$ | 71 |
| Glutaryl | $1.2 \pm 0.02$ | $64.55 \pm 7$ | 309 |

### Effect of different substrate binding at the catalytic site: Docking studies

Despite the interaction of OsCobB with both H3 and H4, it could deacetylate only histone H3 (H3K9Ac and H3K18Ac). It could not deacetylate H4 (tail region) at K5, K12, K16, and K20 positions. [30]. We have tried to explain this discrepancy of H3 and H4 deacetylation by performing the docking studies with HDOCK or Autodock Vina analysis. Molecular docking studies showed the ability of OsCobB to bind H3K9Ac peptide with a binding affinity of −6.4 kcal/mol, which was comparable to that of H4K16Ac (−4.0 kcal/mol) as the receptor is rigid in these cases. Ligplot 2D plots show the catalytic residues involved in interactions with these peptides. Even though both H3K9Ac and H4K16Ac can bind well at the active site, there are differences at the modified lysine site. Generally residues next to the lysine play a big role in binding effectively for catalysis to happen. Next to K16 in H4, there are other longer chain residues which may hamper the entry of peptide at the active site. Probably this question can be better answered by performing MD simulation studies, which will shade light regarding the dynamic nature of peptide after entering the hydrophobic tunnel.

The HDOCK scores for OsCobB-peptide docking are given in **table 3** for comparison. Structurally, all the three OsCobB-peptide complexes overlay really well. There is a kink in the carbon chain of glutaryl group (5 carbon) so that it can interact properly with Tyr55 and Arg58 in a 12Å acyl tunnel. *In-silico* mutagenesis analysis also portrays a significant reduction in the net binding affinity of the peptides on mutation of residues like F23, A25, H110, V151, W152, F153, G154, F183, and P184, some of which are responsible for substrate binding. All these residues are located in the loop regions surrounding the catalytic site and are involved in essential bonded and non-bonded interactions with the lysine modification that facilitate the selectivity of the negatively charged acylations. Additionally, the Y55 and R58 mutations resulted in a drop in the net binding affinity in H3K9 peptides with negative lysine modifications (**Table 4**).

**TABLE 3:** Docking data of different acyl modifications in OsCobB catalytic site.

| **Modified Peptides** | **HDock Docking Score** | **Confidence Score** | **No. of H bonds** | **No. of non-bonded interaction** | **Autodock Vina Binding energy (KJ/mol)** |
| --- | --- | --- | --- | --- | --- |
| H3K9_Ac | -179.9 | 0.65 | 3 | 49 | -26.8 |
| H3K9_mal | -184 | 0.66 | 6 | 55 | -32.2 |
| H3K9_succ | -159 | 0.54 | 7 | 63 | -31.4 |
| H3K9_glut | -169 | 0.6 | 7 | 85 | -28.5 |
| H4K16_Ac | -173 | 0.62 | 6 | 55 | -23.4 |
| ACS_Ac | -154.33 | 0.52 | 5 | 46 | -25.0 |
| IDH2_Ac | -176 | 0.63 | 6 | 58 | -28.0 |

**TABLE 4:** Effects on the binding energy (decrease) by mutating the important residues using *in silico* mutagenesis.

| Modifications | $\Delta\Delta G_{\text{Bind}}$ (kcal/mol) | | | | | | | | |
| --- | --- | --- | --- | --- | --- | --- | --- | --- | --- |
|  | Y55A | R58A | H110G | V151G | W152E | F153G | E155P | Y183A | P184G |
| H3K9_acetyl | 1.5 | 0.3 | 1.45 | 1.65 | 1.9 | 2.02 | 1.73 | 0.63 | 0.68 |
| H3K9_malonyl | 0.9 | 0.7 | 1.97 | 1.8 | 1.5 | 2.4 | 1.6 | 0.04 | 1.74 |
| H3K9_succinyl | 1.64 | 0.65 | 1.7 | 1.53 | 1.6 | 2.5 | 0.6 | 1.1 | 1.35 |
| H3K9_glutaryl | 1.2 | 0.5 | 1.3 | 1.2 | 1.5 | 1.8 | 0.9 | 1.2 | 1.7 |
| H4K16_glutaryl | 1.6 | 0.9 | 1.5 | 1.5 | 1.9 | 1.9 | 0.8 | 0.7 | 1.0 |
| IDH2_Acetyl | 1.5 | 0.5 | 1.8 | 1.36 | 1.1 | 1.9 | 1.6 | 2.2 | 1.0 |
| ACS_Acetyl | 1.3 | 0.7 | 1.1 | 1.4 | 1.2 | 1.5 | 0.8 | 1.9 | 1.5 |

### Effect of OsCobB backbone residues on substrate binding

There is the local structural flexibility in enzyme active site of OsCobB-peptide complexes in comparison to its apo-structure. There are high fluctuations in several loop regions like 19-34, 125-150, 177-184, and 204-208 due to peptide binding. Major movement in the range 125-150 is mainly in the loop region of the OsCobB structure which are involved in substrate binding (FGE). This flexible loop connects the small Zn^2+^ binding domain leading to acyl cavity of the large Rossmann fold domain. Furthermore, mutation studies revealed the relevance of His110, Phe153 in H3K9Acyl peptide binding, whereas Y183 and P184 were also important for binding due to their presence at the entrance of the active site allowing the proper placing of all the peptides. These structural analyses clearly hint at the importance of the protein backbone which may support the effortless binding of the substrate for easy access of the lysine modification for catalysis. The smaller zinc binding domain might have a major role to play in this regard as this domain greatly influences the formation of the catalytic site in all sirtuins.

### Deacylase activity of non-histone targets by OsCobB

Several studies validate the presence of protein acylations in the mitochondria [36]. Similar to lysine acetylation, a significant proportion of succinylated proteins are localized in mitochondria [36,37]. However, there is no proteomic data available regarding the effects of PTMs in ACS and IDH2 in plants which may lead us to pinpoint their effect in cellular metabolism. We have earlier shown that IDH2 and ACS are deacetylated by OsCobB with catalytic efficiency of 422 and 804 M^−1^s^−1^, respectively. Acetylated lysine has a negative effect at the catalytic centre of these enzymes. When the acetyl group was removed by OsCobB, IDH2 enzyme activity increased by atleast 3-fold [34]. Similarly, we wanted to check the ability of OsCobB to remove the other lysine modifications of ACS and IDH2 enzymes. In humans, deacylation of the acidic lysine modifications of ACS and IDH2 leads to variety of disease conditions. We speculate that the outcome of other acidic deacylations by OsCobB will also affect the activities of these enzymes involved in various metabolic networking under different stress conditions.

We found that OsCobB can deacylate acidic modifications in ACS and IDH2 equally well. It follows the same trend as the histones in enzyme efficiency with respect to increase in chain length of the modifications (**Fig 3**). We also reproducibly demonstrated that OsCobB could desuccinylate the mitochondrial substrates with catalytic efficiencies higher than that of lysine deacetylation (**Table 5**). In comparison, its homolog SIRT5 displayed notable desuccinylase and demalonylase activity for ACS with no detectable deacetylation [3]. OsCobB is able to desuccinylate ACS peptide at K642 with comparable efficiency (201 M^−1^s^−1^) as its acetyl counterpart (360 M^−1^s^−1^). This site is homologous to K648 in human ACS, which is a known deacetylation site. Furthermore, SIRT5 desuccinylates IDH2 and thus activating it, facilitating the NADP^+^-dependent oxidative decarboxylation of isocitrate to ***α***-ketoglutarate [7]. So, removal of acidic modifications from ACS or IDH2 will also result in their activations. Interactions between the ligand and OsCobB active site residues are shown in Ligplot figures **(Fig4)**.

**Figure 3:**
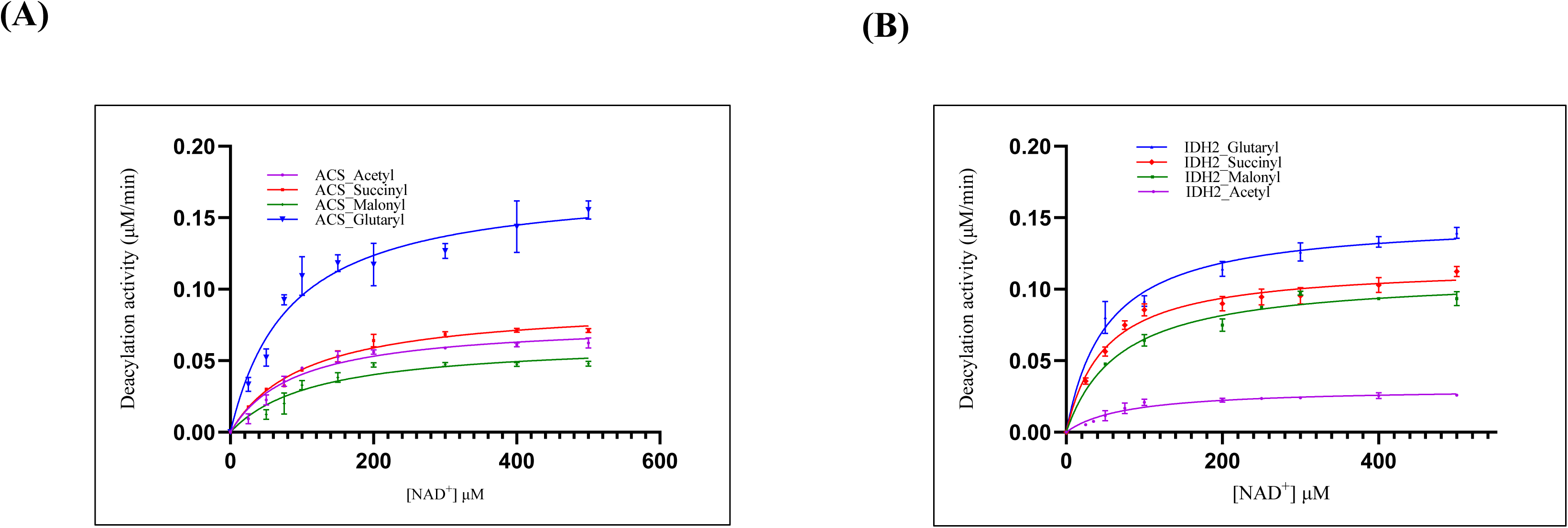
Detection of the preferred deacylase activity of OsCobB with mitochondrial substrate ACS and IDH2. The MM plots of OsCobB for ACS and IDH2 deacylations were calculated using Graphpad Prism 9.1.2 (MM Model). The error bar depicts the S.D.; n=3.

**Figure 4:**
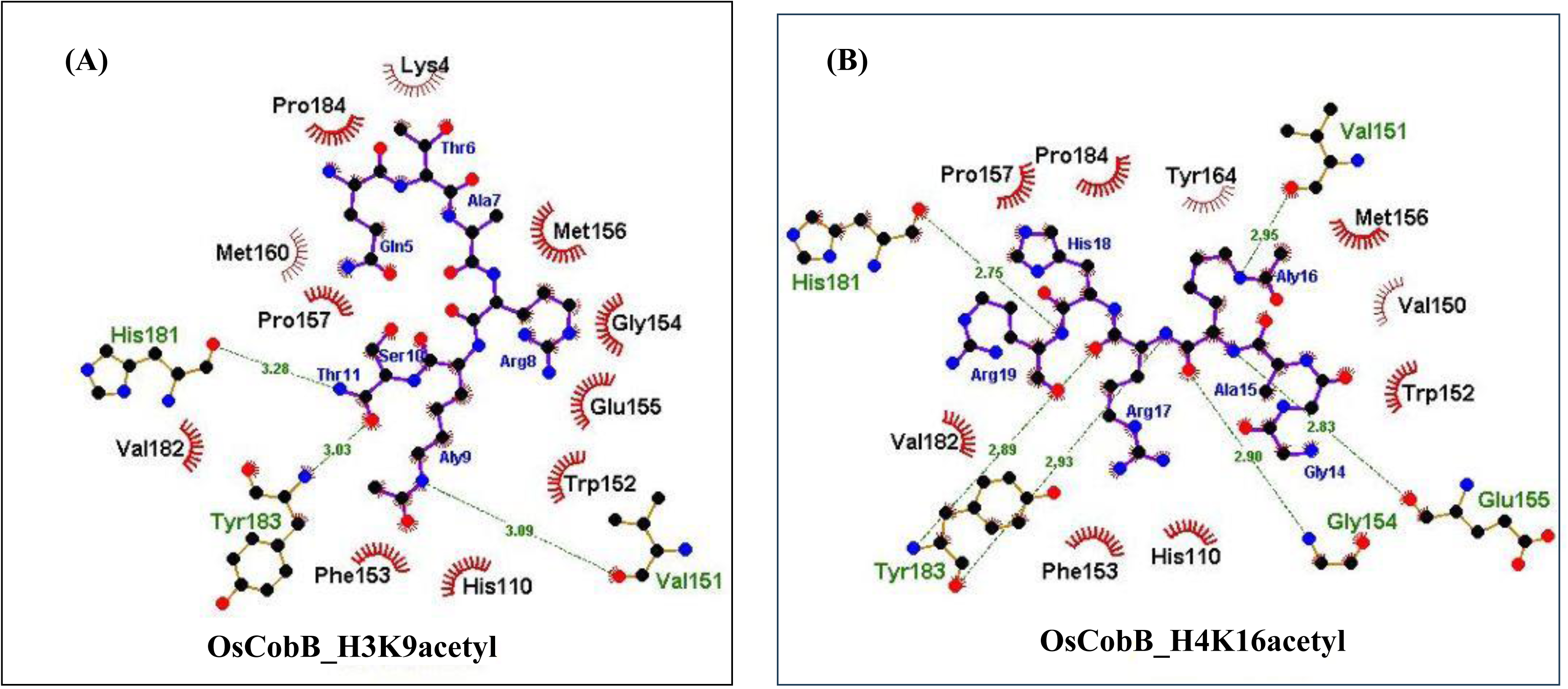
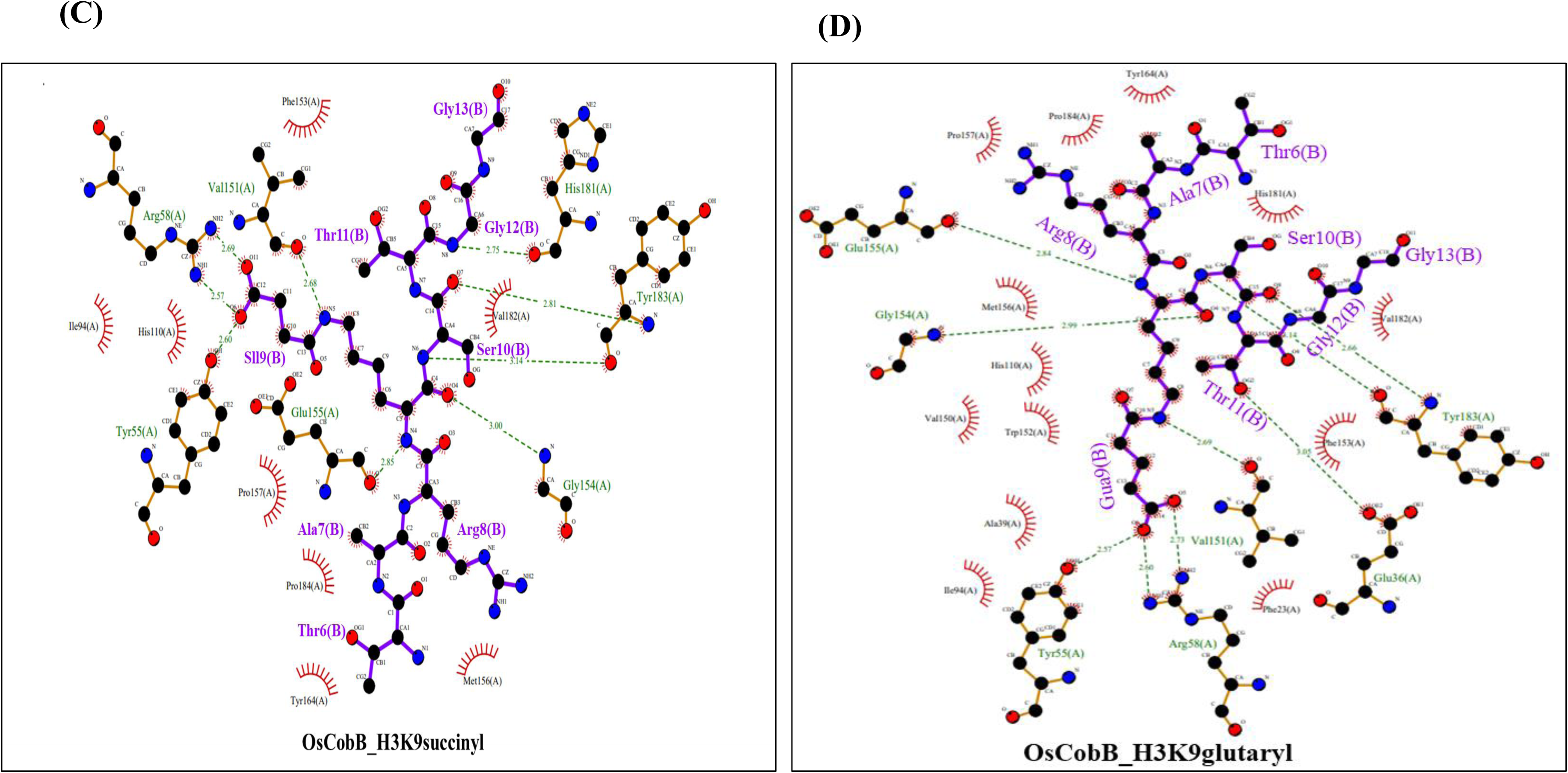
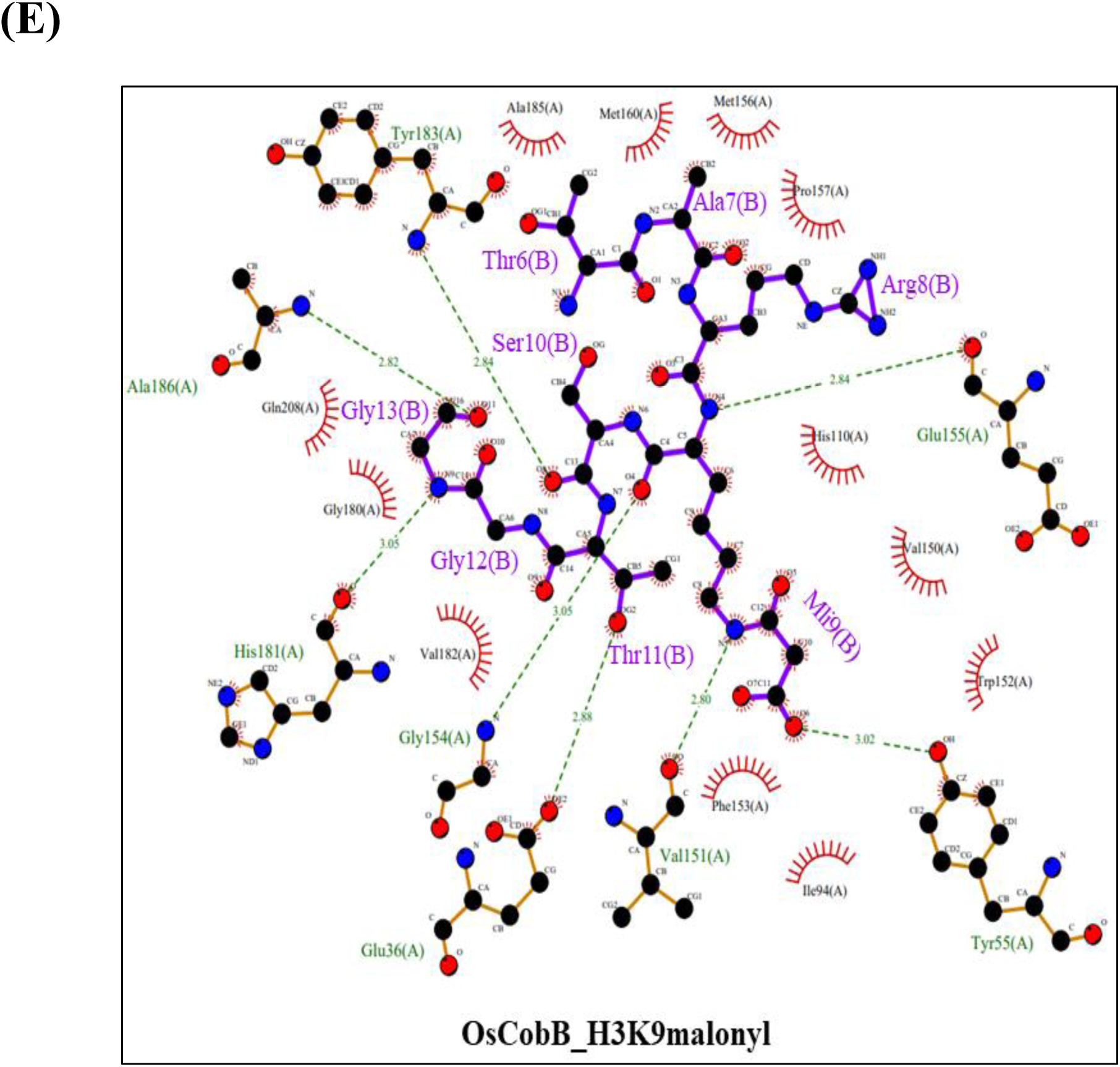
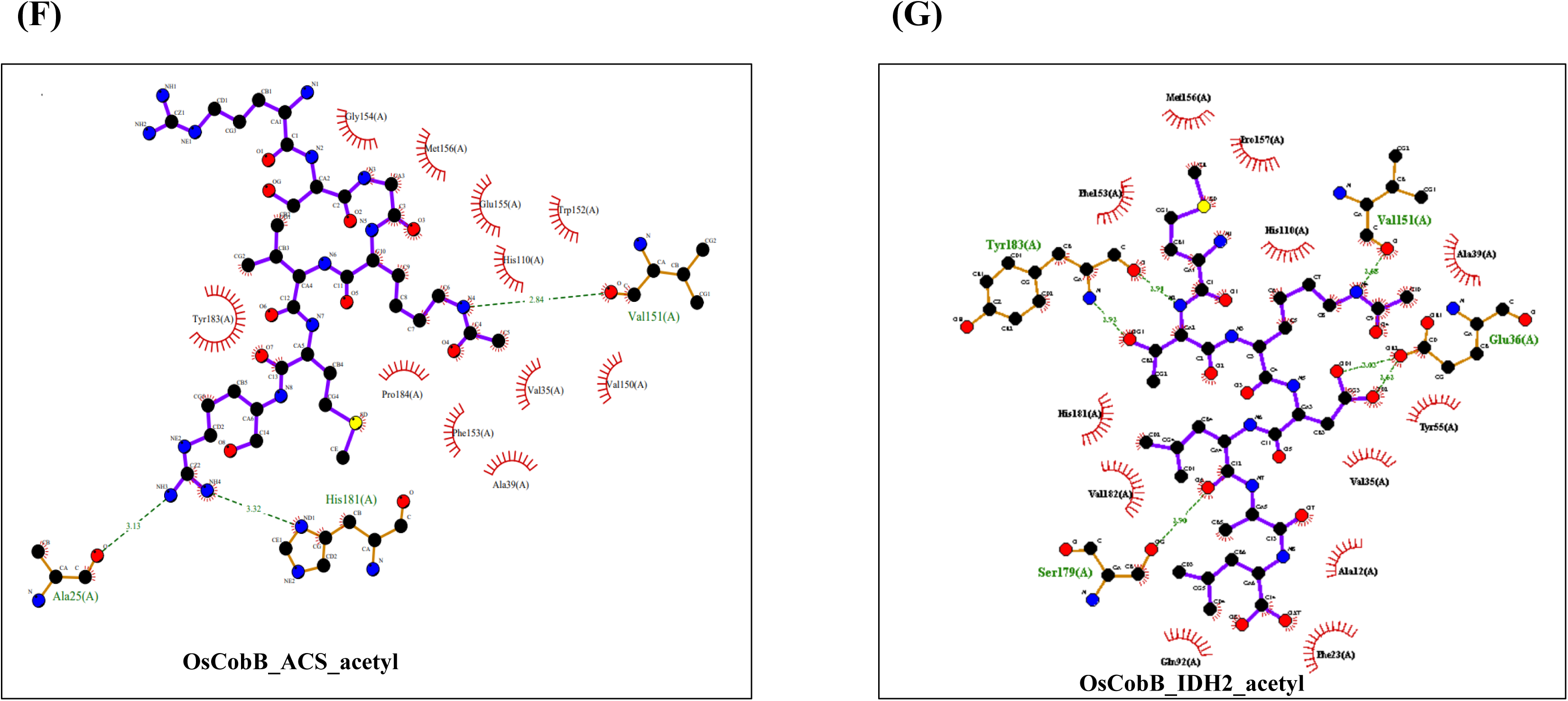
2D interaction Ligplot diagram showing the residues at the OsCobB active site tunnel interacting with the modified lysine peptides.

**TABLE 5:** Comparison of the Michaelis-Menten parameters for NAD^+^ in OsCobB deacylation of IDH2 and ACS obtained from rice leaf extract (mitochondrial fraction)

| **OsCobB**<br>**Substrates** | **$k_{\text{cat}}^{\text{app}}$ ($\text{min}^{-1}$)** | **$K_{\text{m}}^{\text{app}}$ for $\text{NAD}^+$**<br>**($\mu\text{M}$)** | **$k_{\text{cat}}^{\text{app}} / K_{\text{m}}^{\text{app}}$ ($\text{M}^{-1}\text{s}^{-1}$)** |
| --- | --- | --- | --- |
| IDH2_Acetyl | $2.1 \pm 0.13$ | $79.4 \pm 14$ | 441 |
| IDH2_Malonyl | $8.4 \pm 0.35$ | $95 \pm 13.7$ | 1474 |
| IDH2_Succinyl | $7.6 \pm 0.21$ | $46.7 \pm 5$ | 2712 |
| IDH2_Glutaryl | $9.8 \pm 0.4$ | $52 \pm 10$ | 3205 |
| ACS_Acetyl | $5.1 \pm 0.3$ | $89 \pm 14.5$ | 955 |
| ACS_Malonyl | $4.3 \pm 0.4$ | $118.2 \pm 33$ | 607 |
| ACS_Succinyl | $6 \pm 0.2$ | $101.6 \pm 11$ | 984 |
| ACS_Glutaryl | $11 \pm 0.7$ | $65.5 \pm 16$ | 2799 |

### Role of OsCobB-mediated deacylations in stress response

Plants have been shown to regulate their metabolic pathways through multiple PTMs in response to varied stress conditions [38–40]. Till now, histone modifications mostly methylation, acetylation, ubiquitination and phosphorylation have been studied in plants due to their importance in regulation and adaptation due to stress conditions [41]. OsCobB has demonstrated itself as an efficient eraser of different acid modifications on the target proteins. Probably, not all modifications are targeted at the same time, rather depends on the type of environmental pressures the cell encounters [42]. As a result, researchers must investigate the relationship between OsCobB’s ability to deacylate its substrates and its functional mechanism in plants. Taking into account the distinct acylated modifications observed in H3 and H4, this idea was further studied by comparing the degree of histone acylations in rice leaves under various environmental conditions.

Notably, a large number of identified Kac sites are known to overlap with Ksucc/kmal sites, suggesting the existence of substrates which are most likely regulated by more than one PTM [42,43]. Under low temperature and dehydration conditions, the OsCobB expression was found to be 1.5 folds higher in the nucleus than the normal conditions, with subsequent increase also seen in pathogenesis and As toxicity conditions. This hints at the involvement of OsCobB in stress regulation with respect to other histone deacylations. This idea was further explored by comparing the protein expression levels of OsCobB in the nuclear extracts to the degree of acylations in histone H3 and H4 under various stress conditions. An overall reduction in acetylation of H3 was only observed during pathogen attack. Alternately, there was also varying degrees of reduced histone H4 acetylation in all the experimental stress conditions. Pronounced deacetylation of H4 is seen during low temperature and As toxicity, similar to H3 succinylation. However, in times of low temperature, dehydration and arsenic (As) stress, we observed a tremendous reduction in succinylation levels of both H3 and H4. H4 succinylation has also been recorded to assist in DNA repair. There was some reduction in the overall H3 malonyl levels noted during dehydration. Despite the fact that Kmal has been documented to impact protein regulation under drought stress, the proteins involved have not been revealed [44]. Taken together, this could be a crucial step for further exploration of the role of Kmal in environmental stresses. Although OsCobB displayed a great attachment for glutaryl group in both H3 and H4 histones, the stress factors taken into consideration here did not show much alterations. Collectively these data also approve the concept of OsCobB playing a role under varied stress stimuli displaying quantifiable deacylase activities, though their accurate regulation mechanism is not clear. (**Fig 5**)

**Figure 5:**
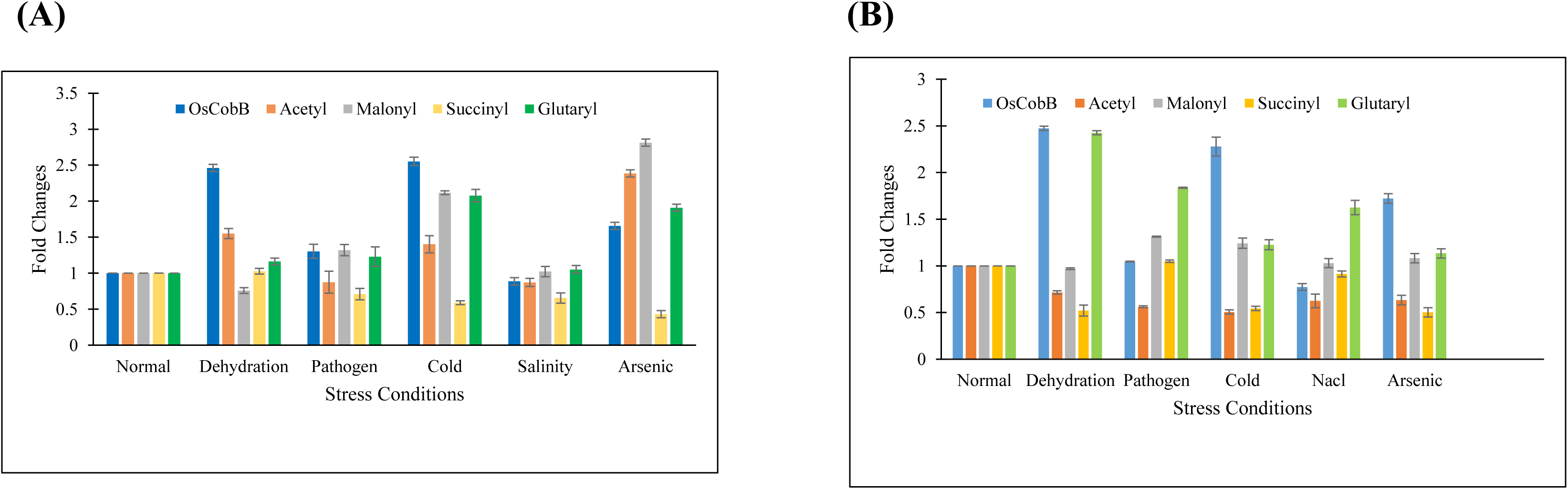
Detection of lysine deacylation levels in endogenous histones (A) H3 and (B) H4 related to OsCobB protein expression under different stress conditions.

Since OsCobB is capable of removing both acetyl and succinyl moiety from IDH2, it is possible that the selectivity of its acylation may be triggered by different stress stimuli. Previous experiments could correlate the 1.5-fold increase in OsCobB protein expression in the mitochondria to high salinity, low temperature stress and pathogenesis [30]. Besides, a 2-fold rise in IDH2 deacetylation under low temperature stress was distinguishable from the 5-fold rise in IDH2 desuccinylation under pathogen stress. IDH2 deacetylation is an important step in plant TCA cycle for energy production in mitochondria. This enzyme is also involved in NADP^+^ homeostasis. Previous study has demonstrated that Lys391 in OsIDH2, which is preserved as Lys413 in human IDH2, is the site of deacetylation that increases its activity. This site is critical for NADP^+^ binding and thus affects its activity. OsCobB was also found to be involved in the deacetylation and further activation of IDH2. SIRT3 is in charge of deacetylating IDH2, mostly during antioxidant defence [45]. Disruptions of SIRT5 gene results in hyper-succinylation of IDH2 leading to cancer progression. Similarly, we investigated the regulation of OsCobB activity under stress because we were intrigued by its ability to deacylate IDH2 very efficiently. Our findings showed numerous environmental factors like dehydration, cold, salinity, iron and cyanide toxicity, that were linked to increased OsCobB protein expression in the mitochondria. With no impact on its acetylation, we saw a significant drop in IDH2_Succ in response to pathogen infection and cyanide exposure in the plants. In contrast, during iron toxicity in the soil, IDH2Ac levels were drastically reduced. We have also seen reduced succinyl level in ACS during cold and cyanide stress (**Fig 6**). It will be beneficial to further clarify the metabolic pathways and variables involved behind these actions, as a significant amount of this information is comparatively limited in plants. Plants are often facing different soil conditions with various metal toxicity. How they are able to cope with these toxicities and make modifications in their system to survive, needs to be seen.

**Figure 6:**
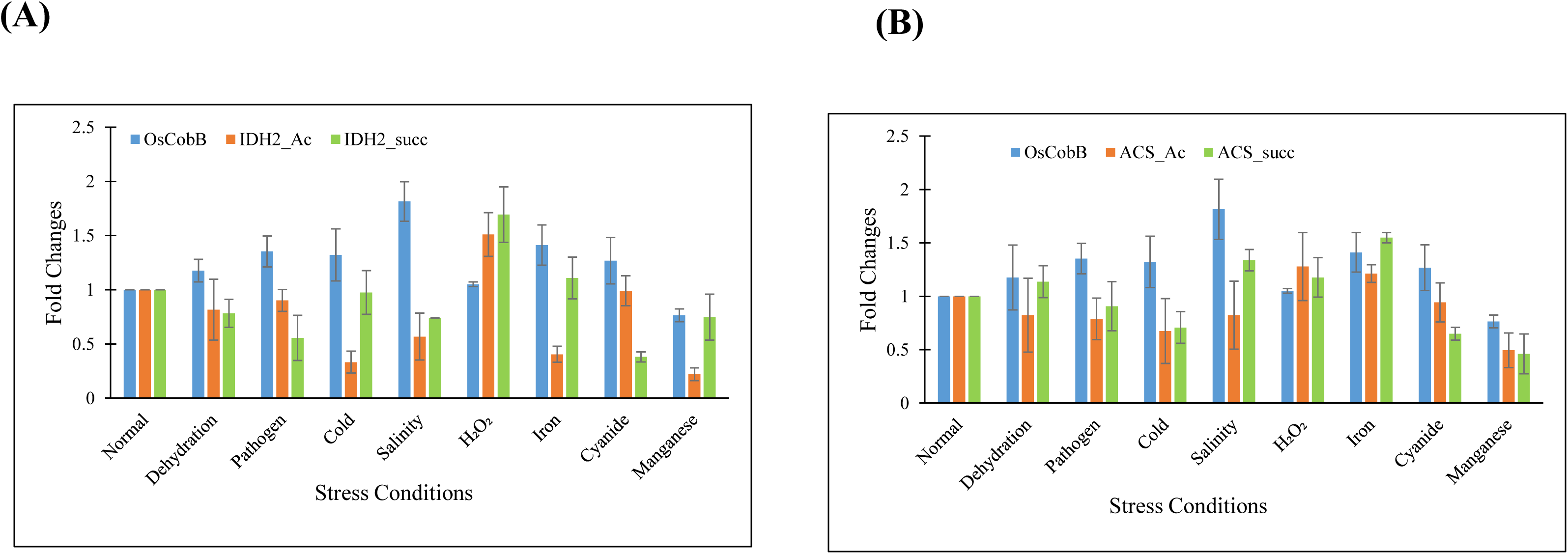
Comparative analysis of deacylated mitochondrial substrates (ACS and IDH2) in relation to OsCobB expression under various stress conditions.

### Conclusions

There are very limited studies available regarding the role of these deacylation reactions in plants and how it is affecting the plant physiology. The presence of acetyl and some of these acidic lysine modifications (malonyl, succinyl) has been shown in plants like tomato, rice seeds, pepper, patchouli plant leaves using western blot and mass spectrometry [42,43,46,47]. No proteomic studies on the glutarylation of plant proteins is available to understand the metabolic networking of this modification. We have detected the presence of negatively charged acyl modifications in nucleus as well as mitochondria of the rice leaf extract by using the anti-Pan acyl immunoblots. Preferred sites for the acidic modifications in histones were discussed. We have also studied the deacylations of mitochondrial proteins ACS and IDH2, which have not been reported earlier in plants. Thus, we have analyzed an eraser which is capable of making changes to these modifications under different stress conditions.

The molecular basis of OsCobB action was elucidated using structural and biochemical analysis. Docking analysis showed that OsCobB can accommodate a diverse range of lysine modifications in its active site (hydrophobic tunnel), with a preference for acetyl, glutaryl, succinyl, and malonyl lysine side chains, respectively. This was due to the presence of the conserved catalytic site residues (Tyr55 and Arg58), which interacts with the acidic modifications of the lysine side chain. However, not every interacting ligand could be deacylated at every potential lysine position, as clearly shown in this study. This is due to both the peptide backbone and its proper orientation, important in selecting the target for modification. Also, the placement of other catalytically important residues like His110, Phe153, Phe183, Pro184 lining the peptide binding site mattered. Class III sirtuins preferred acidic modifications due to the presence of conserved Tyr and Arg at the active site but SIRT7 which is not a member of class III also favoured desuccinylation. This suggests that there are also other structural factors in sirtuins responsible for this selectivity.

The acidic modifications are similar in structure with increase in one -CH_2_ moiety and known to regulate variety of proteins in different pathways. If these negatively charged modifications are introduced in the middle of the protein, it can affect the structure and folds of the protein which will eventually affect its function, protein-protein interactions. We have taken into consideration several stress conditions like drought, high salinity, cold, pathogen, As toxicity etc. which affects the plant growth and development, and eventually agricultural production. We have further tried to relate the extent of changes in the acidic modifications with respect to OsCobB protein expressions. This is the first instance to document all of these changes in acidic lysine modifications in rice leaves under varying plant stress situations. H3K9 deacetylation was found to be connected to H4 desuccinylation and deacetylation, which protect against drought and cold tolerance. As toxicity also caused a decrease in both H4 Ac and Succ levels, which targeted OsCobB activity in response to the stress. The overexpression of the OsCobB protein under these stress circumstances suggests a role of this enzyme in removal of these histone modifications. Depending on the stress conditions, the sirtuin played a varied role in different subcellular compartments. It was found to be overexpressed in the nucleus during dehydration, arsenic toxicity and cold stress, and in the mitochondria during pathogenesis, Iron toxicity and high salinity stress. Mitochondria is considered as a hub for plant growth and development as they house several different enzymes which can be modified. This provides preliminary data for future studies on plant stress situations in connection to ACS/IDH2 lysine deacylation. IDH2 Ac/Succ levels fluctuated a lot with respect to OsCobB expression in pathogen, cold and metal toxicity. As we know that deacylations of these enzymes activates them, their relevance in this case cannot be ignored.

There is a correlation between the different PTM levels with various stress conditions. Also in humans, variations in these deacylation levels in various tissues lead to different disease conditions. The diversity of acyl incorporation may have a role in modulating protein-macromolecule interactions under various stress conditions, ultimately influencing metabolic control. This calls for more in-depth research into OsCobB-like proteins in other plants and their regulations. It is critical to investigate the factors causing co-localization and varied protein expression during stress. There is also a possibility of crosstalk between different modifications to tackle the stress situation in plants. As it is still unclear how these modifications affect the plant during various stress and who are the key players involved in this.

The comprehensive analysis of these reversible PTMs will support the future studies of protein modifications in the field of enhanced crop productivity. There are several examples of other classes of HDACs linked with plant stress conditions. OsCobB may also influence additional growth and developmental pathways in plants. Due to its open active site architecture which allows access to most of the modifications, it needs to be seen what other acyl group modifications OsCobB is capable of erasing.

## Abbreviations

PTM: post translational modification,
HDAC: histone deacetylase,
NAD^+^: nicotinamide adenine dinucleotide,
ACS: acetyl CoA synthetase,
IDH2: isocitrate dehydrogenase 2

## Acknowledgement

This work was funded by Department of Science and Technology, Govt of India (CRG/2019/003037).

